# Taste cues elicit prolonged modulation of feeding behavior in *Drosophila*

**DOI:** 10.1101/2022.05.27.493715

**Authors:** Julia U. Deere, Anita V. Devineni

## Abstract

Taste cues regulate immediate feeding behavior, but their ability to modulate future behavior has been less well-studied. Pairing one taste with another can modulate subsequent feeding responses through associative learning, but this requires simultaneous exposure to both stimuli. We investigated whether exposure to one taste modulates future responses to other tastes even when they do not overlap in time. Using *Drosophila*, we found that brief exposure to sugar enhanced future feeding responses, whereas bitter exposure suppressed them. This modulation relies on neural pathways distinct from those that acutely regulate feeding or mediate learning-dependent changes. Sensory neuron activity was required not only during initial taste exposure but also afterward, suggesting that ongoing sensory activity may maintain experience-dependent changes in downstream circuits. Thus, the brain stores a memory of each taste stimulus after it disappears, enabling animals to integrate information as they sequentially sample different taste cues that signal local food quality.

## INTRODUCTION

The taste system evolved to regulate feeding behavior, ensuring that animals consume caloric foods and avoid potential toxins. Different types of tastes, such as sweet or bitter, generally activate separate populations of sensory cells to drive or suppress behavioral responses related to feeding (Liman et al., 2014; Scott, 2018). In addition to regulating immediate feeding behavior, taste cues can also modulate future responses through learning. For example, studies in *Drosophila* have shown that pairing sugar with either natural or optogenetically-triggered bitter taste diminishes subsequent feeding responses to sugar (Kirkhart and Scott, 2015; Masek et al., 2015; Jelen et al., 2021). Pairing taste cues with odors can also modify future odor responses (Schwaerzel et al., 2003; Krashes et al., 2009; Huetteroth et al., 2015; Das et al., 2014). Learned taste-taste and odor-taste associations in *Drosophila* are mediated by the mushroom body, the primary learning and memory center of the fly brain (Kirkhart and Scott, 2015; Masek et al., 2015; Jelen et al., 2021; Schwaerzel et al., 2003; Krashes et al., 2009; Huetteroth et al., 2015; Das et al., 2014).

These taste learning paradigms, in which one taste modulates the future response to another taste, have used simultaneous or near-simultaneous exposure to both taste cues (Kirkhart and Scott, 2015; Masek et al., 2015; Jelen et al., 2021). In a natural environment, animals may encounter different taste cues sequentially rather than simultaneously. Does the experience of one taste modulate an animal’s subsequent response to a different taste? Since taste cues signal the local food quality, It would be adaptive for animals to integrate taste information over time as they make feeding decisions. Exposure to an appetitive taste such as sugar may enhance the likelihood of future feeding, whereas aversive tastes such as bitter may suppress it. Indeed, the ability of sugar to sensitize subsequent feeding responses to water was suggested by early studies in insects (Dethier et al., 1965; Duerr and Quinn, 1982).

In this study, we investigated whether exposure to one taste modulates future responses to other tastes even when they do not overlap in time. We found that brief exposure to sugar enhanced future feeding responses over seconds to minutes, whereas bitter exposure suppressed these responses. Optogenetic activation of taste sensory neurons recapitulated this behavioral modulation. Interestingly, the activity of sensory neurons was required not only during initial taste exposure but also afterward, suggesting an ongoing role for sensory activity in maintaining a taste memory. Neuronal manipulations show that this prolonged modulation relies on neural pathways distinct from those that acutely regulate feeding or mediate learning-dependent changes in feeding behavior. These findings reveal a new form of behavioral flexibility in the taste system, enabling cross-modulation between different taste pathways even when they are not simultaneously active.

## RESULTS

### Brief exposure to sweet taste enhances future feeding responses

We began by asking whether brief taste exposure affects the likelihood of future feeding-related responses in *Drosophila*. We focused on the proboscis extension response (PER), which represents the initiation of feeding, as a proxy for feeding behavior (Dethier, 1976; Scott, 2018). We hypothesized that sugar exposure would enhance the likelihood of appetitive taste responses such as PER, whereas bitter exposure would decrease this likelihood. We first tested whether sugar exposure affects subsequent PER to water (Figure 1A), which normally induces very low levels of PER in water-satiated flies. The percent of flies showing PER to water dramatically increased when tested 20 seconds after a brief (∼1 second) exposure to sugar on the proboscis (Figure 1B). 7% of flies showed PER to water before sugar exposure, whereas 46% showed PER after sugar exposure (Figure 1B). For all subsequent analyses, we quantify this modulatory effect as the difference in PER: (% PER after sugar exposure) – (% PER before sugar exposure).

**Figure 1:**
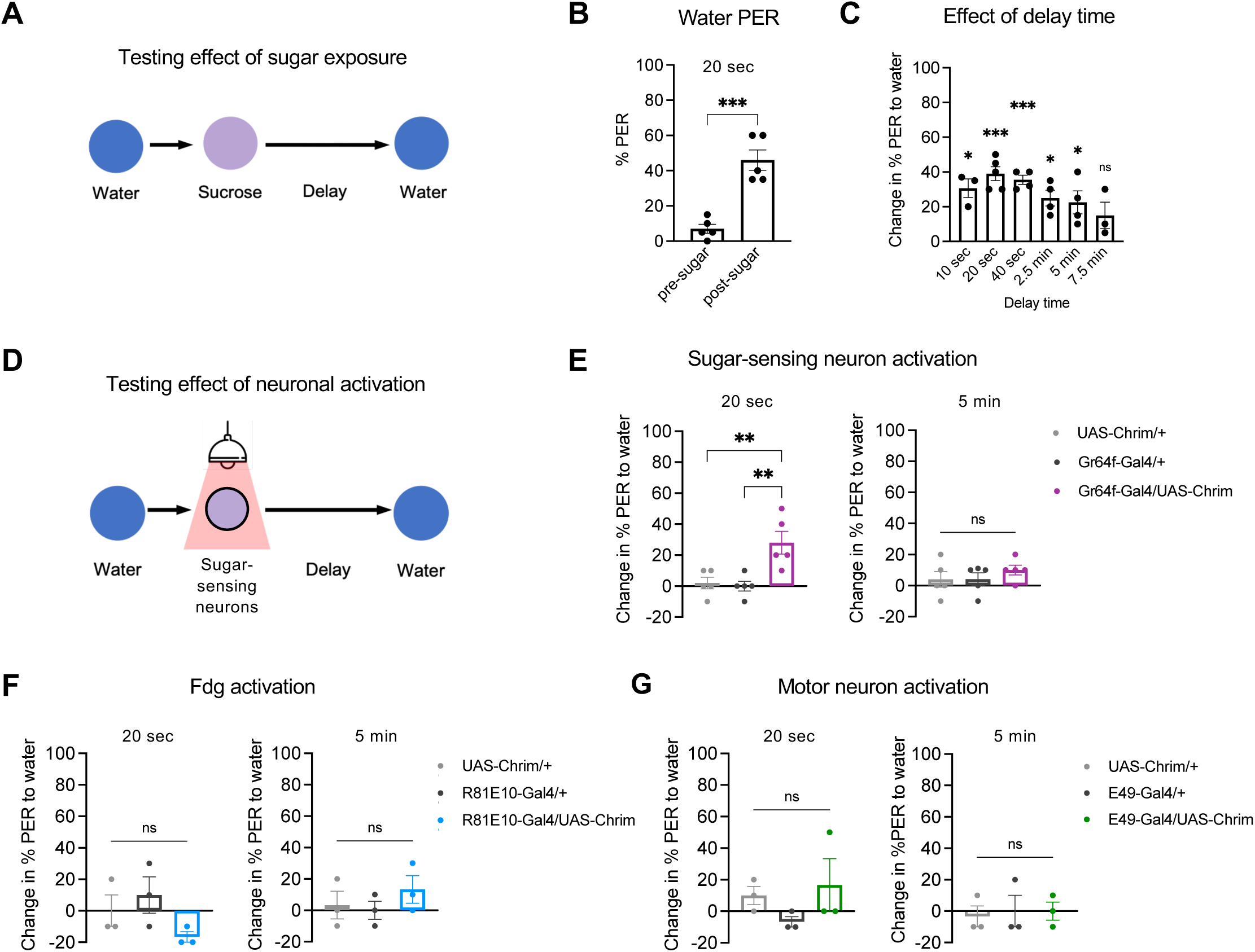
Brief sugar exposure increases the likelihood of PER to water over short timescales. (A) Schematic of experiment design to test the effect of brief (∼1 sec) sugar exposure on subsequent responses to water. (B) Percent of flies showing PER to water before and 20 sec after sugar presentation (n = 5 sets of flies, paired t-test). (C) Change in the percent of flies showing PER to water before and after sugar exposure for different delay periods (n = 3-5 sets of flies). The change is calculated as a difference, and positive values indicate increased PER after sugar exposure. Values were compared to zero using one-sample t-tests. (D) Schematic of experiments using optogenetic stimulation of sugar-sensing neurons. Light was delivered for ∼1 sec. For all optogenetic experiments, experimental flies carrying both the *Gal4* and *UAS* transgenes were compared to control flies carrying only one of the two transgenes. (E-G) Change in PER to water 20 sec (left panels) or 5 min (right panels) after optogenetic activation of sugar-sensing neurons (E; n = 5 sets of flies), the Fdg neuron (F; n = 3 sets of flies), or the MN9 proboscis motor neuron (G; n = 3 sets of flies). *Gr64f-Gal4* (E), *R81E10-Gal4* (F), or *E49-Gal4* (G) was used to drive *UAS-Chrimson*. Genotypes were compared using one-way ANOVA followed by Tukey’s post-tests. For all figures: Error bars represent standard error of the mean. Data points represent values for each set of flies (∼10 or 20 flies; see Methods). *p<0.05, **p<0.01, ***p<0.001, ****p<0.0001, ns = not significant.

To determine how long this modulation lasts, we tested various delay times between sugar exposure and the subsequent water stimulus. PER to water increased most substantially (30-40% higher than baseline) when tested within 10 to 40 seconds of sugar exposure, after which the effect gradually decreased (Figure 1C). By 7.5 minutes, PER to water was statistically indistinguishable from baseline levels (Figure 1C). For all subsequent experiments, we used a 20 second delay to test PER modulation and a 5 minute delay as the control condition in which modulation has typically decayed to near zero.

We wanted to confirm that the increase in PER to water after sugar exposure was not due to residual sugar remaining on the proboscis. We therefore replaced sugar exposure with optogenetic activation of sweet-sensing neurons (Figure 1D). We used *Gr64f-Gal4* to drive *UAS-Chrimson*, which encodes a light-gated cation channel (Klapoetke et al., 2014). Experimental flies were compared to control flies carrying only one of the two transgenes. Light stimulation was delivered for ∼1 second, analogous to sugar exposure in the previous experiments. In an earlier study we showed that optogenetic activation of taste sensory neurons using Chrimson does not elicit prolonged neuronal activity after the light is turned off (Devineni et al., 2021). In the experimental flies, PER to water increased by ∼30% when tested 20 seconds after optogenetic activation of sugar-sensing neurons, but did not show a significant increase after a 5 minute delay (Figure 1E). Thus, sweet taste enhances PER to a different stimulus over short timescales, and this effect is due to transient activation of sweet-sensing neurons rather than residual sugar on the proboscis.

We next asked whether this modulatory effect on PER could also be elicited by activating downstream neurons in the sugar pathway. We tested optogenetic activation of the Fdg “feeding command” neuron (Flood et al., 2013) or the MN9 proboscis motor neuron (Gordon and Scott, 2009; Schwarz et al., 2017), both of which reside downstream of sugar-sensing neurons. We confirmed that activation of both neurons acutely elicited PER (Figure S1), as previously reported (Flood et al., 2013; Gordon and Scott, 2009; Schwarz et al., 2017). However, neither Fdg nor MN9 activation enhanced PER to water 20 seconds later (Figure 1F-G), showing that they are not sufficient to elicit prolonged PER modulation. These results imply that different neural pathways downstream of sugar-sensing neurons are responsible for acutely regulating PER versus modulating the likelihood of future PER.

### Brief exposure to bitter taste decreases the likelihood of future feeding responses

Given that appetitive taste elicited a prolonged increase in the likelihood of PER, we asked whether the opposite type of modulation could be observed with an aversive taste. We tested whether brief (∼1 second) exposure to bitter taste modulates PER to sugar tested 20 seconds later (Figure 2A). Two bitter compounds, quinine and lobeline, strongly suppressed PER to sugar 20 seconds later, with the percent of responding flies decreasing from ∼90% to ∼30% for quinine (Figure 2B) and from ∼70% to ∼20% for lobeline (Figure 2C). When PER to sugar was tested 5 minutes after bitter exposure, significant PER suppression was still observed with lobeline, but not quinine (Figure 2B-C). We used quinine as the bitter stimulus for subsequent experiments.

**Figure 2:**
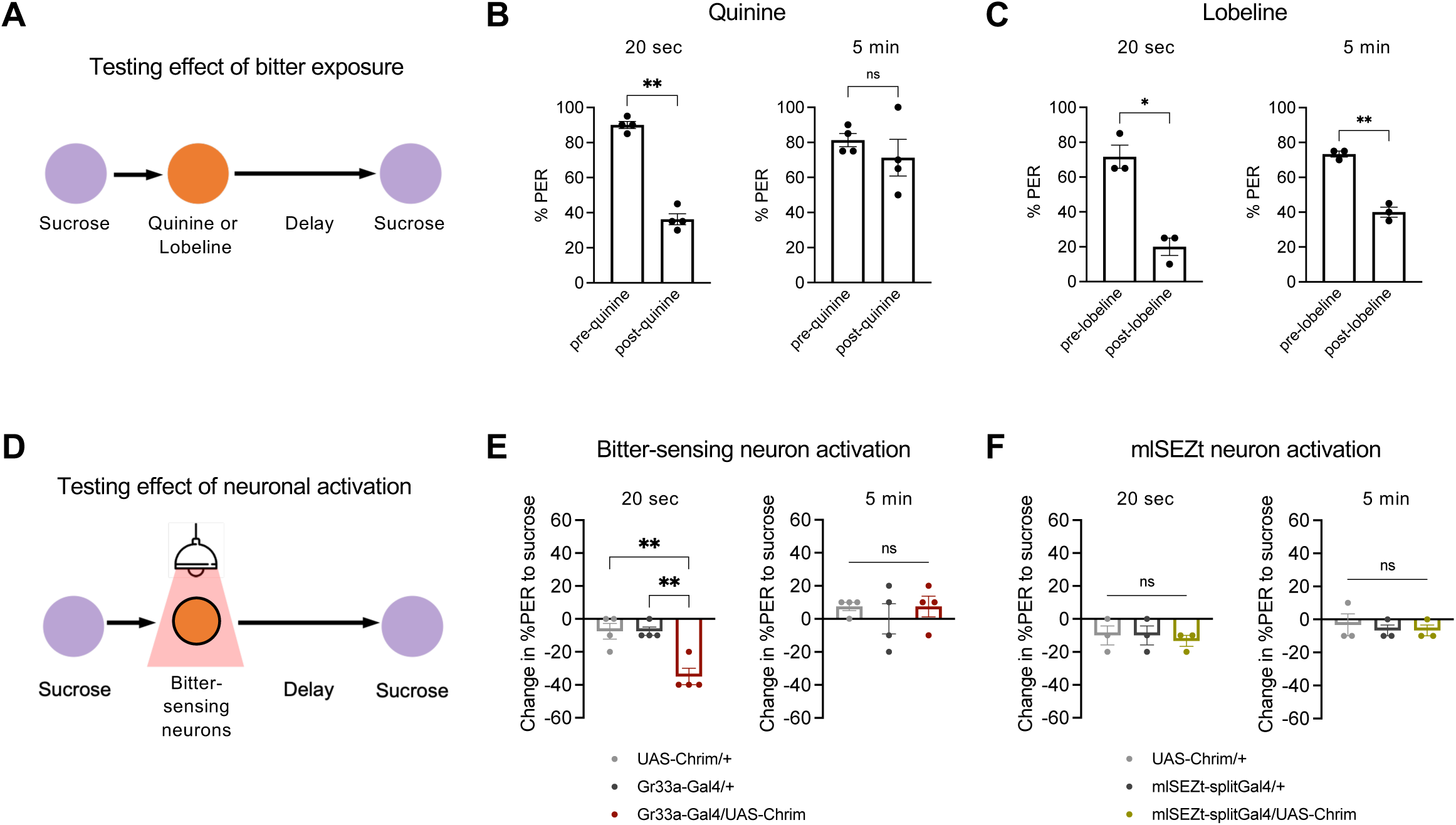
Brief exposure to bitter taste decreases the likelihood of subsequent PER to sugar. (A) Schematic of experiment design to test the effect of brief (∼1 sec) bitter exposure on subsequent responses to sugar. (B-C) Percent of flies showing PER to 100 mM sucrose before and after exposure to 10 mM quinine (B; n = 4 sets of flies) or 5 mM lobeline (C; n = 3 sets of flies) (paired t-tests). Left panels represent a 20 sec delay between the bitter and sugar stimuli; right panels represent a 5 min delay. (D) Schematic of experiments using optogenetic stimulation of bitter-sensing neurons. Light was delivered for ∼1 sec. (E-F) Change in PER to sucrose tested 20 sec (left panels) or 5 min (right panels) after activation of bitter-sensing neurons (E; n = 4 sets of flies) or mlSEZt second-order bitter neurons (F; n = 3 sets of flies). *Gr33a-Gal4* (E) or *mlSEZt-splitGal4* (*R29F12-AD* + *R55E01-DBD*) (F) was used to drive *UAS-Chrimson*. Genotypes were compared using one-way ANOVA followed by Tukey’s post-tests.

We then tested whether optogenetic activation of bitter-sensing neurons, labeled by *Gr33a-Gal4*, could recapitulate this prolonged suppression of PER (Figure 2D). Brief activation of bitter neurons elicited a significant decrease in PER to sugar when tested 20 seconds later, but not after 5 minutes (Figure 2E). Together, these results show that prolonged modulation of PER is bidirectional and can be driven by multiple taste modalities, with sugar enhancing and bitter suppressing the likelihood of future PER.

We recently identified a set of second-order bitter neurons, termed mlSEZt neurons, that receive input from bitter-sensing cells and acutely suppress PER to sugar when activated (Deere et al., 2022). We therefore tested whether mlSEZt activation could also suppress future PER to sugar. PER to sugar tested 20 seconds after light stimulation was similar to baseline PER and did not differ between experimental and control flies (Figure 2F). Similar to our manipulations of downstream sugar neurons (Figure 1F-G), these results show that downstream taste neurons that acutely regulate PER do not necessarily elicit prolonged modulation of PER. Thus, parallel pathways downstream of taste sensory neurons modulate PER on different timescales.

### Sensory neuron activity is required during and after taste stimulation for prolonged PER modulation

We next asked whether the activity of taste sensory neurons is required for prolonged modulation of PER by sweet or bitter tastes. We silenced sugar-or bitter-sensing neurons using *UAS-GtACR*, which encodes a light-gated chloride channel (Figure 3A-F) (Mohammad et al., 2017). As expected, silencing sugar-or bitter-sensing neurons during the sugar or bitter exposure, respectively, reduced or abolished the prolonged PER modulation elicited by these stimuli (Figure 3B, 3E). We then tested the effect of silencing sensory neurons during the delay period between the sugar or bitter stimulus and the test stimulus (water or sugar). Unexpectedly, neuronal silencing during the delay period reduced the modulatory effect of sugar or bitter to a similar degree as silencing during the sugar or bitter stimulus itself (Figure 3C, 3F). These results suggest that the activity of sensory neurons is required not only during the initial taste exposure but also afterward.

**Figure 3:**
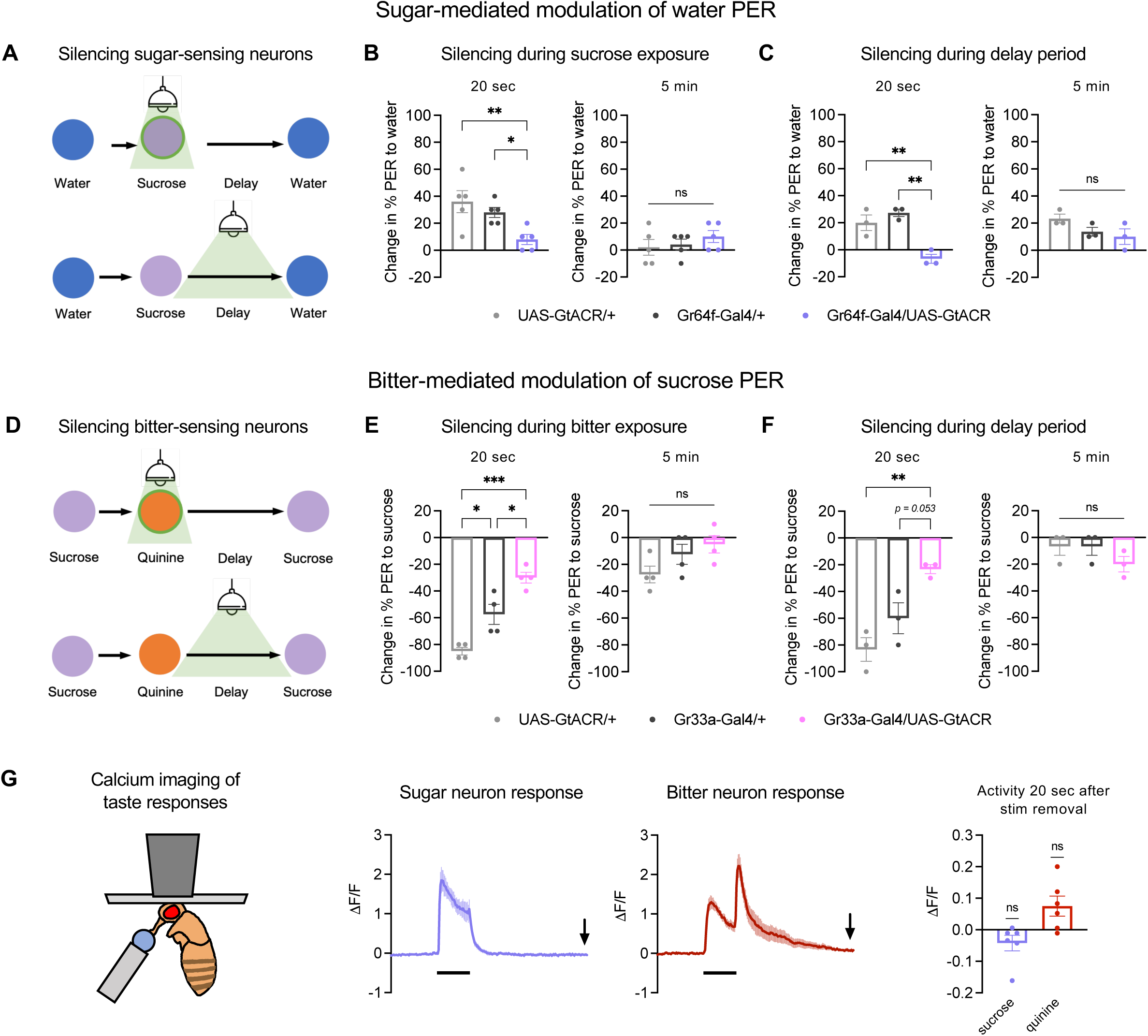
Sensory neuron activity is required during and after taste stimulation for prolonged PER modulation. (A) Schematic of experiment design to optogenetically silence sugar-sensing neurons during or after the sugar stimulus. (B-C) Change in PER to water tested 20 sec (left panels) or 5 min (right panels) after sugar exposure. Sugar-sensing neurons were silenced during the sugar stimulus (B) or during the delay period between the sugar and bitter stimuli (C). n = 5 sets of flies. *Gr64f-Gal4* was used to drive *UAS-GtACR1*. (D) Schematic of experiment design to optogenetically silence bitter-sensing neurons during or after the bitter stimulus. (E-F) Change in PER to sugar tested 20 sec (left panels) or 5 min (right panels) after bitter exposure. Bitter-sensing neurons were silenced during the bitter stimulus (E, n = 4 sets of flies) or during the delay period between the bitter and sugar stimuli (F, n = 3 sets of flies). *Gr33a-Gal4* was used to drive *UAS-GtACR1*. (G) Left: Schematic of setup to image sensory neuron activity in response to taste stimulation on the proboscis. The wings and legs were taped behind the fly and are not shown. Middle: GCaMP traces showing activity of sugar-or bitter-sensing neurons during and after stimulation with sucrose or quinine, respectively. Black bars indicate duration of stimulus (5 sec), and arrows indicate the time point 20 sec after stimulus removal. Right: Quantification of GCaMP activity 20 sec after taste removal (n = 6 flies, one-sample t-test comparing responses to zero, which represents the pre-stimulus baseline). *Gr64f-Gal4* (sugar neurons) or *Gr33a-Gal4* (bitter neurons) was used to drive *UAS-GCaMP6f*. In panels B-F, genotypes were compared using one-way ANOVA followed by Tukey’s multiple comparisons test.

The effect of sensory neuron silencing during the delay period was surprising because taste sensory neurons are not thought to show prolonged activity following stimulus removal. In our own previous studies, the only type of post-stimulus activity we observed was a transient offset response in bitter-sensing neurons (Devineni et al., 2021; Devineni et al., 2019). To more carefully examine sensory neuron activity during a longer post-stimulus period, we performed two-photon calcium imaging of sugar-and bitter-sensing neurons using GCaMP6f (Chen et al., 2013). We imaged the axon terminals of these neurons, located in the subesophageal zone (SEZ) of the brain, while delivering taste stimuli (sucrose or quinine) to the proboscis (Figure 3G). We observed stimulus-locked responses consistent with those observed previously (Devineni et al., 2021). Sugar and bitter neurons responded to the onset of sucrose or quinine, respectively, and bitter neurons also responded to quinine offset (Figure 3G). Following stimulus removal, sugar neuron activity returned to baseline within a few seconds, whereas bitter neurons first showed an offset response and returned to baseline more slowly (Figure 3G). By 20 seconds after the sugar or bitter stimulus, the time point at which a strong modulatory effect on PER is observed (Figures 1 and 2), sugar and bitter neuron activity had returned to baseline (Figure 3G). These results show that although silencing sensory neuron activity after sugar or bitter exposure strongly reduces their modulatory effect tested 20 seconds later, sensory neurons are not activated above baseline levels at this time. Thus, their baseline activity may be important for maintaining changes in downstream circuits that store the memory of the taste stimulus.

### Bitter activation does not modulate subsequent responses of sugar-sensing neurons

Our behavioral results show that activation of sugar-or bitter-sensing neurons modulates future PER to other tastes (Figures 1 and 2), implying that the activation of one taste modality elicits changes in the activity of neural circuits processing other taste modalities. For bitter exposure, the subsequent suppression of PER to sugar suggests that one or more nodes of the sugar circuit become less responsive to sugar input. We asked whether this modulation may occur at the level of sugar sensory neurons, which have been shown to exhibit state-dependent modulation (Inagaki et al., 2012). We imaged the sucrose response of sugar-sensing neurons 20 seconds after optogenetic activation of bitter neurons, thus recapitulating our behavioral paradigm. While PER to sugar was strongly suppressed with this protocol (Figure 2), the responses of sugar-sensing neurons were not affected (Figure 4). These results suggest that bitter exposure suppresses subsequent PER to sugar by modulating sugar responses downstream of sugar-sensing neurons.

**Figure 4:**
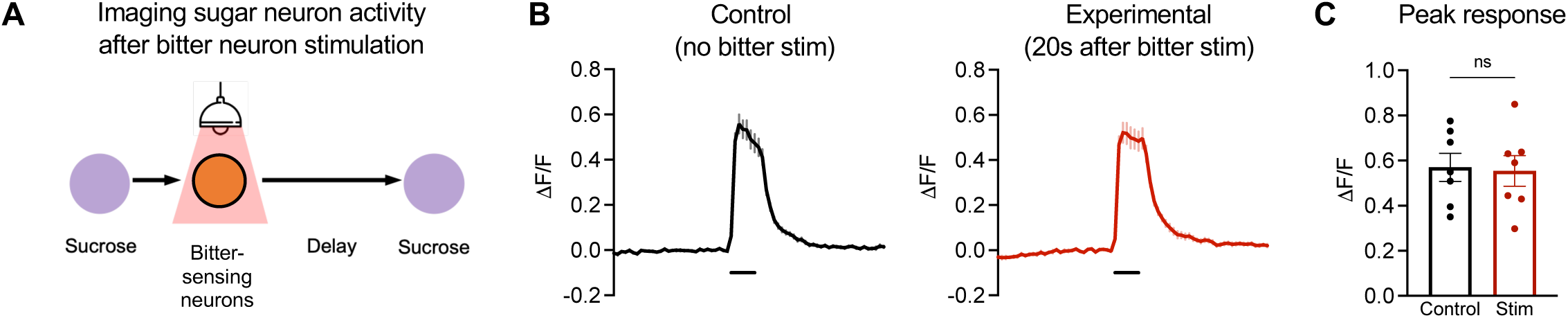
Bitter activation does not modulate subsequent responses of sugar-sensing neurons. (A) Schematic of experiment protocol. We recorded the responses of sugar-sensing neurons to a sugar stimulus presented 20 sec after optogenetic activation of bitter neurons. Flies carried *Gr66a-lexA* driving *lexAop-Chrimson* and *Gr64f-Gal4* driving *UAS-GCaMP6f*. (B) GCaMP traces for experimental trials (red), in which sugar neurons were imaged 20 sec after bitter neuron activation, and control trials (black), which did not include bitter stimulation (n = 7 flies, 3-5 trials each). (C) Peak response during the sugar stimulus for each group (unpaired t-test). Dots represent averages for individual flies.

### Mushroom body circuits are not required for prolonged PER modulation

The imaging results described above reveal that, in our taste modulation paradigm, sugar and bitter tastes do not interact at the level of sensory neurons. These tastes must therefore interact within downstream taste circuits. The mushroom body receives sugar and bitter input and mediates learning-dependent interactions between sugar and bitter taste (Kirkhart and Scott, 2015; Modi et al., 2020; Liu et al., 2012; Devineni et al., 2021; Masek et al., 2015). When a sugar stimulus is paired with bitter taste, PER to sugar is subsequently suppressed (Kirkhart and Scott, 2015; Masek et al., 2015). This form of associative learning is thought to involve co-activation of taste-responsive Kenyon cells, the intrinsic neurons of the mushroom body, with modulatory dopaminergic neurons (DANs), resulting in plasticity of Kenyon cell synapses onto mushroom body output neurons (Modi et al., 2020). We tested whether the same circuit mediates the non-associative modulation of PER in our paradigm. We first silenced Kenyon cells by expressing Kir2.1, an inwardly rectifying potassium channel (Baines et al., 2001). These flies still showed prolonged modulation of PER elicited by either sugar or bitter, similar to controls: sugar exposure increased subsequent PER to water (Figure 5A), and bitter exposure decreased subsequent PER to sugar (Figure 5B).

**Figure 5:**
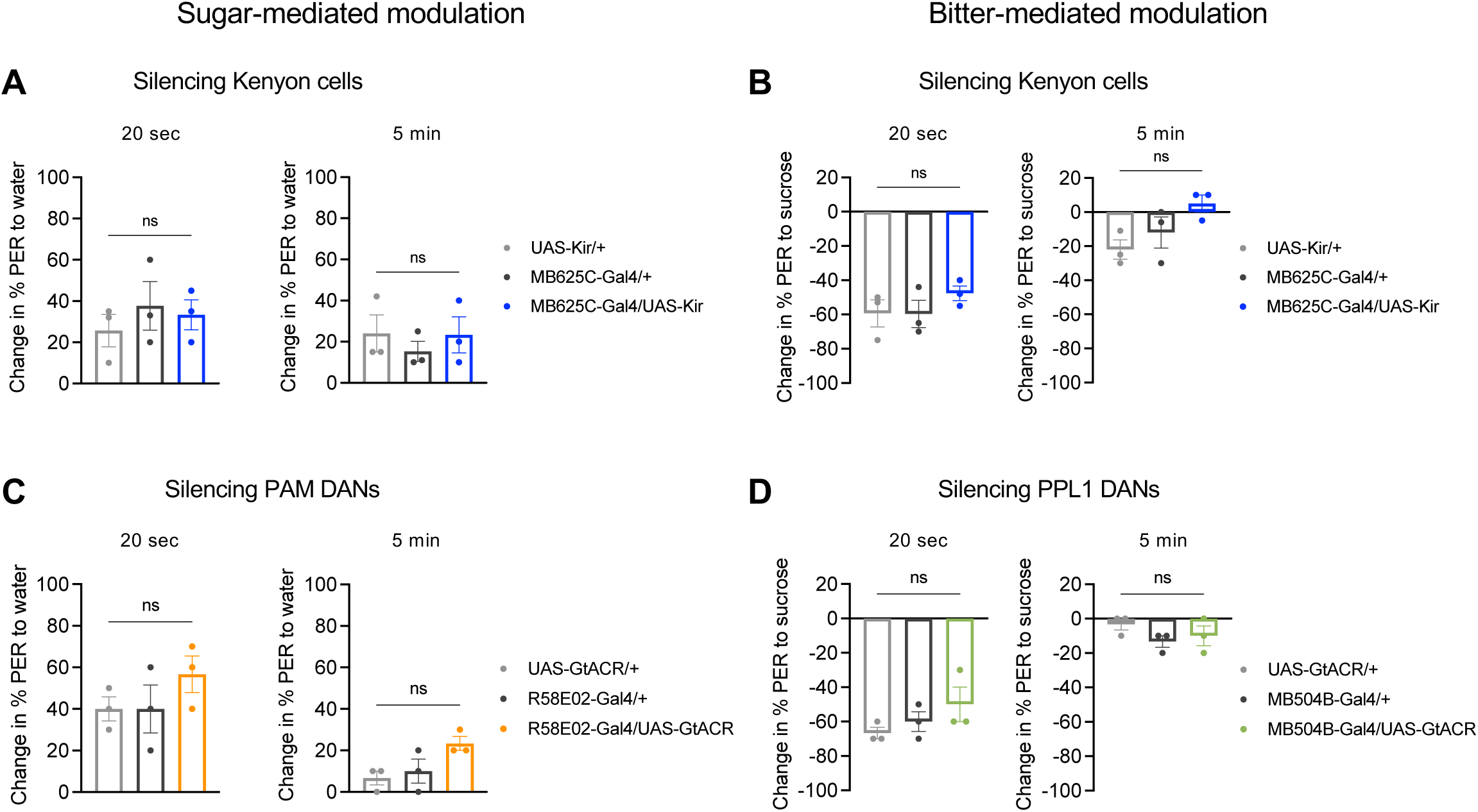
Mushroom body circuits are not required for prolonged PER modulation. (A-B) Effect of constitutively silencing Kenyon cells (*MB0625C* split-*Gal4* driving *UAS-Kir2*.*1*) on PER modulation induced by sugar (A) or bitter (B). (A) Change in PER to water tested 20 sec (left) or 5 min (right) after sugar exposure (n = 3 sets of flies). (B) Change in PER to sucrose tested 20 sec (left) or 5 min (right) after quinine exposure (n = 3 sets of flies). (C) Effect of optogenetically silencing PAM DANs (*R58E02-Gal4* driving *UAS-GtACR1*) on PER modulation by sugar. Graphs show change in PER to water tested 20 sec (left) or 5 min (right) after sugar presentation. Green light was turned on during the sugar stimulus. (D) Effect of optogenetically silencing PPL1 DANs (*MB504B* split*-Gal4* driving *UAS-GtACR1*) on PER modulation by bitter taste. Graphs show change in PER to sucrose tested 20 sec (left) or 5 min (right) after quinine presentation. Green light was turned on during and after the bitter stimulus, but was turned off before the sugar stimulus. In all panels, genotypes were compared using one-way ANOVA followed by Tukey’s multiple comparisons test.

We also silenced subsets of DANs innervating the mushroom body. Protocerebral anterior medial (PAM) DANs are activated by appetitive stimuli such as sugar, whereas protocerebral posterior lateral cluster 1 (PPL1) DANs respond to aversive stimuli such as bitter (Modi et al., 2020; Liu et al., 2012; Kirkhart and Scott, 2015; Devineni et al., 2021). Optogenetic silencing of PAM DANs with GtACR during sugar exposure did not affect the ability of sugar to enhance subsequent PER to water (Figure 5C). Similarly, optogenetic silencing of PPL1 DANs during bitter exposure did not affect the ability of bitter taste to reduce subsequent PER to sugar (Figure 5D). Taken together, these results suggest that mushroom body circuits are not required for the prolonged, non-associative modulation of PER in our paradigm.

## DISCUSSION

In this study, we characterized a new type of behavioral modulation in the taste system. Brief exposure to an appetitive or aversive taste bidirectionally modulates the likelihood of future feeding responses to other stimuli over seconds to minutes. In a natural environment, this modulation would enable an animal to integrate information over time as it samples different food sources. Encountering an appetitive or aversive taste likely signals that the local environment contains food sources of high or low quality, respectively, allowing the animal to adjust its likelihood of feeding based on recent history.

We found that optogenetic activation of taste sensory neurons recapitulated the prolonged behavioral modulation. Interestingly, the activity of sensory neurons was required not only during taste exposure but also afterward, suggesting that ongoing sensory activity may maintain experience-dependent changes in downstream circuits. Calcium imaging revealed that bitter modulation of PER to sugar occurs downstream of sensory neurons. Neuronal manipulations suggest that prolonged PER modulation relies on pathways distinct from those that acutely regulate PER or mediate learning-dependent changes in PER.

Together, these findings reveal several new insights into the organization and function of taste circuits. First, neural pathways for processing distinct taste modalities are capable of modulating one another even when they are not simultaneously active. Second, the brain can store the memory of a taste stimulus after it disappears, potentially through persistently active neurons or synaptic changes (see below), and post-stimulus sensory neuron activity is critical for maintaining this memory. Third, the sugar and bitter taste pathways each branch into multiple circuits that regulate the same feeding response in different contexts, revealing a division of labor between parallel pathways. These results establish a paradigm for future work to further dissect the cellular and circuit mechanisms underlying prolonged behavioral modulation by taste.

### Circuit mechanisms for prolonged PER modulation

Most studies have investigated cross-modulation between different taste modalities when they are presented at overlapping times. Different tastes presented together are acutely integrated to determine behavior (Wang et al., 2004; Inagaki et al., 2014) and can modulate future responses through associative learning (Kirkhart and Scott, 2015; Masek et al., 2015; Jelen et al., 2021). Because behavioral modulation in our paradigm does not require simultaneous exposure to both tastes, the underlying circuit mechanisms are likely to be distinct from those previously described. Indeed, we found that prolonged PER modulation in our paradigm does not require the mushroom body, which mediates associative taste learning (Kirkhart and Scott, 2015; Masek et al., 2015; Jelen et al., 2021).

We propose a possible circuit mechanism underlying prolonged modulation of PER by sugar and bitter (Figure 6). First, the activation of sugar or bitter sensory neurons causes a memory of the taste stimulus to be stored in the downstream circuit. By “memory”, we refer to any type of neural change that encodes the previous taste experience after the taste has disappeared. This memory then modulates activity in other taste pathways when they are activated. The memory of sugar modulates activity in the water-sensing pathway, and the memory of bitter modulates activity in the sugar-sensing pathway.

**Figure 6:**
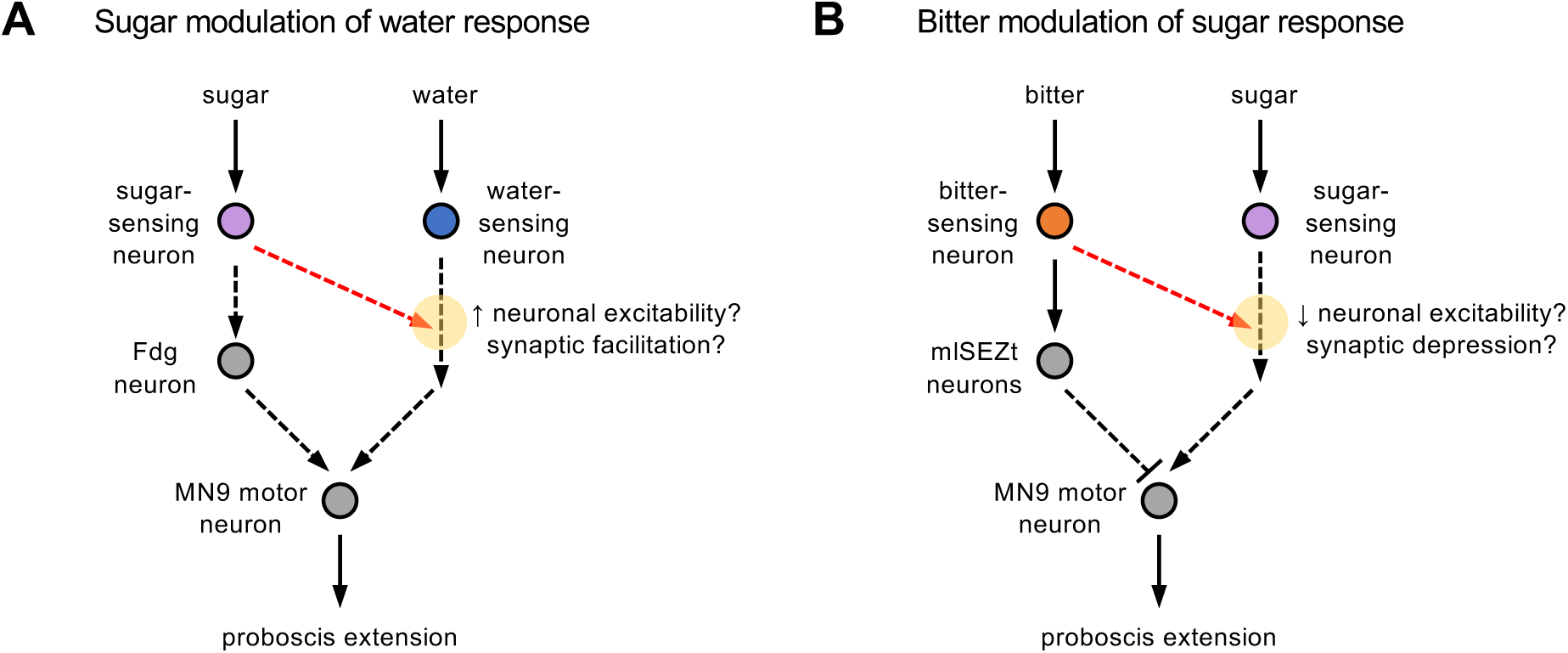
Circuit model for prolonged modulation of PER across different taste modalities. Models for prolonged modulation of PER by sugar (A) or bitter (B). Dotted arrows indicate connections that may be indirect. Red arrows indicate downstream branches of the sugar (A) or bitter (B) pathways that elicit prolonged modulation of responses to other tastes (water or sugar, respectively). Yellow circles denote sites of modulation in the water (A) or sugar (B) pathways following previous sugar or bitter exposure, respectively. We propose that the memory of previous taste exposure could be stored as persistent activity in the modulatory pathway (red arrows) or as neuronal or synaptic plasticity at the convergence site between the two pathways (yellow circles).

Separate memories may be stored for sugar and bitter tastes within their respective circuits. Alternatively, the brain may store a general memory of recent taste experience that has either an appetitive or aversive valence. In either case, it is possible that this memory is encoded as a short-lived internal state that modulates a variety of behaviors beyond feeding. Several studies have described internal states that are induced by brief exposure to a sensory stimulus. In flies, this includes states corresponding to locomotor arousal (Lebestky et al., 2009), defensive arousal or “fear” (Gibson et al., 2015), and sexual arousal (Jung et al., 2020; Hindmarsh Sten et al., 2021). A chemosensory stimulus, carbon dioxide, has been shown to elicit a persistent state of blood feeding in female mosquitoes (Sorrells et al., 2022). Persistent responses to brief sugar exposure have been also been described, in which flies show local search behavior in an attempt to locate the previous sugar stimulus (Corfas et al., 2019; Behbahani et al., 2021). These studies show that flies can store a memory of the sugar stimulus over short timescales, but they differ from our study in that they describe persistent behavioral changes, whereas we characterize the ability of sugar to modulate future stimulus responses without eliciting overt behavioral changes in the absence of a stimulus.

How is the memory of a taste stimulus encoded in our paradigm? Given the short-lived nature of this memory, it is unlikely to involve canonical plasticity mechanisms such as changes in receptor trafficking or gene expression (Mayford et al., 2012). Instead, it may be mediated by mechanisms proposed to underlie working memory, which include persistent neuronal activity and short-term changes in synaptic strength (Zylberberg and Strowbridge, 2017; Barak and Tsodyks, 2014). For example, the activation of taste sensory neurons could elicit persistent, slowly decaying activity in downstream neurons. Persistent activity in the sugar pathway would transmit excitation to the water-responsive circuit, enhancing PER to a future water stimulus, whereas persistent activity in the bitter pathway would transmit inhibition to the sugar circuit, suppressing PER to a future sugar stimulus (Figure 6). As in other systems, persistent activity could be generated through intrinsic cellular properties or recurrent connections (Zylberberg and Strowbridge, 2017). Alternatively, prolonged PER modulation could involve changes in synaptic strength. The modulated synapses would need to reside at or downstream of convergence nodes between different taste pathways. Sugar exposure could induce synaptic facilitation at a downstream synapse that is also activated by water, whereas bitter exposure could induce synaptic depression at a downstream synapse that is normally activated by sugar (Figure 6).

Either model must account for the observation that silencing sensory neurons following the taste stimulus strongly reduces the behavioral modulation (Figure 3A-F) even though sensory neurons are not activated above baseline for most of this period (Figure 3G). One possibility is that the mechanism underlying plasticity – whether a persistently active neuron or synaptic change – integrates sensory activity in a continuous and bidirectional manner. For example, persistently active neurons would be activated by brief sensory neuron activation, but the resulting persistent activity can be reduced if sensory neuron activity is then silenced below baseline. A previous study reported a behavioral effect of GtACR silencing of sugar-sensing neurons in the absence of sugar (Mohammad et al., 2017), supporting the idea that baseline activity of sugar neurons is read out by downstream circuits.

### Parallel pathways for regulating a single feeding response

The downstream neurons that mediate prolonged PER modulation are apparently distinct from known neurons that acutely regulate PER, such as Fdg and mlSEZt neurons. The pathways for acute and prolonged PER modulation are also distinct from the circuit that mediates PER through associative learning, the mushroom body. Thus, the sugar and bitter pathways diverge into multiple branches that regulate PER on different timescales. This division of labor to modulate a single behavioral output, PER, may shed light on why the processing of a single taste involves multiple parallel pathways. Recent anatomical studies have revealed that pathways for individual taste modalities diverge into several branches at the first synapse (Talay et al., 2017; Snell et al., 2020; Deere et al., 2022). We have proposed that these parallel pathways may regulate different behaviors (Deere et al., 2022), but the present results also suggest that they may modulate the same behavior, PER, in different contexts.

### Sites of modulation in the taste system

In our PER modulation paradigm, the site where the initial taste memory is stored may be distinct from the site where subsequent taste behaviors are modulated. For example, persistent activity in downstream bitter-sensing neurons may be transmitted as feed-forward inhibition onto the sugar-sensing circuit (Figure 6). In this case the neurons receiving the inhibition would represent the site of modulation. Our imaging experiments suggest that this modulation does not occur at the level of sensory neurons. Although several studies have observed modulation of taste sensory neurons, such as by hunger or dietary experience (Inagaki et al., 2012; Inagaki et al., 2014; LeDue et al., 2016; May et al., 2019; Wang et al., 2020; Ganguly et al., 2021), modulation also occurs in downstream taste circuits (Kain and Dahanukar, 2015; Devineni et al., 2019; Shiu et al., 2022). It will be interesting to determine whether the prolonged PER modulation that we have characterized occurs at the same site as other types of modulation, suggesting the presence of core modulatory nodes for modulating taste behavior.

## ACKNOWLEDGEMENTS

We thank Richard Axel for his generous support; Barbara Noro for general advice, manuscript comments, and sharing unpublished fly lines; Chris Rodgers for feedback on the manuscript; and the Janelia Research Center and the Bloomington Drosophila Stock Center (BDSC) for providing fly strains.

## AUTHOR CONTRIBUTIONS

A.V.D conceived and supervised the project. J.U.D. conducted all behavioral experiments and A.V.D. conducted all imaging experiments. J.U.D. and A.V.D analyzed data, generated figures, and wrote the manuscript.

## STAR METHODS RESOURCE AVAILABILITY

### Lead contact

Further information and requests for resources and reagents should be directed to and will be fulfilled by the lead contact, Anita Devineni.

### Materials availability

This study did not generate new unique reagents.

### Data and code availability

- All data reported in this paper will be shared upon request to the lead contact.
- Imaging analysis was performed using existing code; references are cited in the Key Resources Table and Methods.
- Any additional information required to reanalyze the data reported in this paper is available from the lead contact upon request.

## EXPERIMENTAL MODEL AND SUBJECT DETAILS

### Fly strains and maintenance

Flies were reared at 25ºC on standard cornmeal food. Experiments were performed on mated females. Behavioral experiments were performed on 3-7 day-old flies, while calcium imaging was performed on 1-2 week-old flies to ensure robust GCaMP6f expression. Flies used for optogenetic experiments were maintained in constant darkness throughout development and housed on food containing 1 mM all trans-retinal for 2-4 days prior to testing. Flies used for behavioral testing or imaging during the behavioral paradigm (Figure 4) were food-deprived with water for one day. For optogenetic experiments, 1 mM all trans-retinal was added to the water during food deprivation.

*2U* was used as the wild-type control strain. Genotypes used for each experiment are specified in the figures and legends, and source information for fly strains is provided in the Key Resources Table.

## METHOD DETAILS

### Behavioral assays

PER assays were conducted using previously described methods (Devineni et al., 2019). Briefly, flies were anesthetized on ice, immobilized on their backs with myristic acid, and the two anterior pairs of legs were glued down so that the proboscis was accessible for taste stimulation. Flies recovered from gluing for 30-60 minutes in a humidified chamber. Flies were water-satiated before testing. To test PER, tastants were briefly applied to the proboscis (∼1 second) using a small piece of Kimwipe. The same procedure was used for taste exposure to elicit modulation of subsequent PER. Tastants used were water, 100 mM sucrose, 10 mM quinine, or 5 mM lobeline. To activate taste neurons with Chrimson, a red LED (617 nm) was manually turned on for 1 second. To silence neurons with GTACR, a green LED (530 nm) was manually turned on during the specified portion of the experiment. LEDs were controlled by an Arduino.

For PER, flies were tested sequentially in groups of ∼10 for optogenetic experiments and ∼20 for non-optogenetic experiments. Experimental and control flies were tested on the same days. For non-optogenetic experiments, flies in the same group were glued onto a single glass slide. For optogenetic experiments, each fly was glued onto a separate dish so that the light stimulus would not affect other flies. In each case, the percent of flies showing PER to each stimulus was manually recorded. Only full proboscis extensions were counted as PER. At the end of each experiment, flies were tested with a positive control (500 mM sucrose) and were excluded from analysis if they did not respond. For statistical analyses, each set of ∼10 or 20 flies was considered to be a single data point (“n”). At least 3 independent sets of flies were tested for each experiment, representing a minimum of 30 flies per experiment.

### Calcium imaging

Flies were imaged using previously described protocols and equipment (Devineni et al., 2019; Devineni et al., 2021). Briefly, flies were immobilized with tape and the proboscis was secured in an extended position. An imaging window was cut on the anterior surface of the head, the antennae were removed, and the esophagus was cut. Tastants were delivered and removed using a custom-built solenoid pinch-valve system controlled by MATLAB software. Proper taste delivery was monitored using a side-mounted camera. Imaging experiments were performed using a two-photon laser scanning microscope (Ultima, Bruker) equipped with an ultra-fast Ti:S laser (Chameleon Vision, Coherent). The excitation wavelength was 925 nm, and emitted photons were collected with a GaAsP photodiode detector through a 60X water-immersion objective. A single plane through the most dense area of axonal projections was chosen for imaging.

## QUANTIFICATION AND STATISTICAL ANALYSIS

### Calcium imaging analysis

Calcium imaging data were analyzed using MATLAB code from previous studies (Devineni et al., 2019; Devineni et al., 2021). Images were registered within and across trials to correct for movement in the x-y plane using a sub-pixel registration algorithm. Regions of interest (ROIs) were drawn manually. Average pixel intensity within the ROI was calculated for each frame. The average signal for 20 frames preceding stimulus delivery was used as the baseline signal, and the ΔF/F (change in fluorescence divided by baseline) for each frame was then calculated. The peak response was quantified as the average ΔF/F value for the two highest consecutive frames during stimulus presentation. Trials were excluded if the tastant drop failed to make proper contact with the proboscis based on video monitoring.

### Statistical testing

Statistical analyses were performed using GraphPad Prism, Version 9. Statistical tests and results are reported in the figure legends. We used one-sample t-tests to compare individual groups to an expected value (e.g. zero), paired or unpaired t-tests to compare two groups, and one-way ANOVA with Tukey’s post-tests to compare more than two groups. All graphs represent mean ± SEM. Sample sizes are listed in the figure legends.

## KEY RESOURCES TABLE

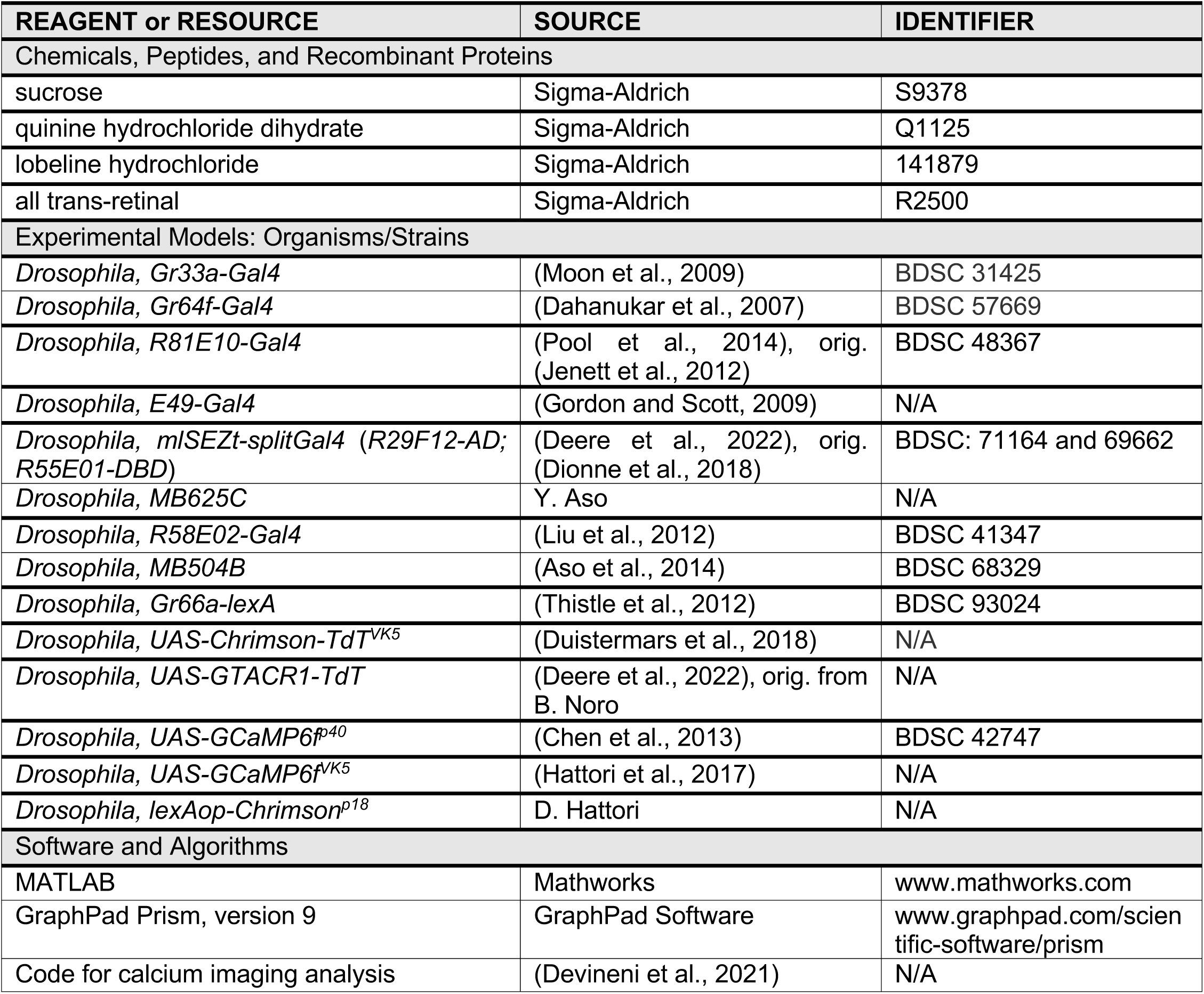

---

**Figure S1:**
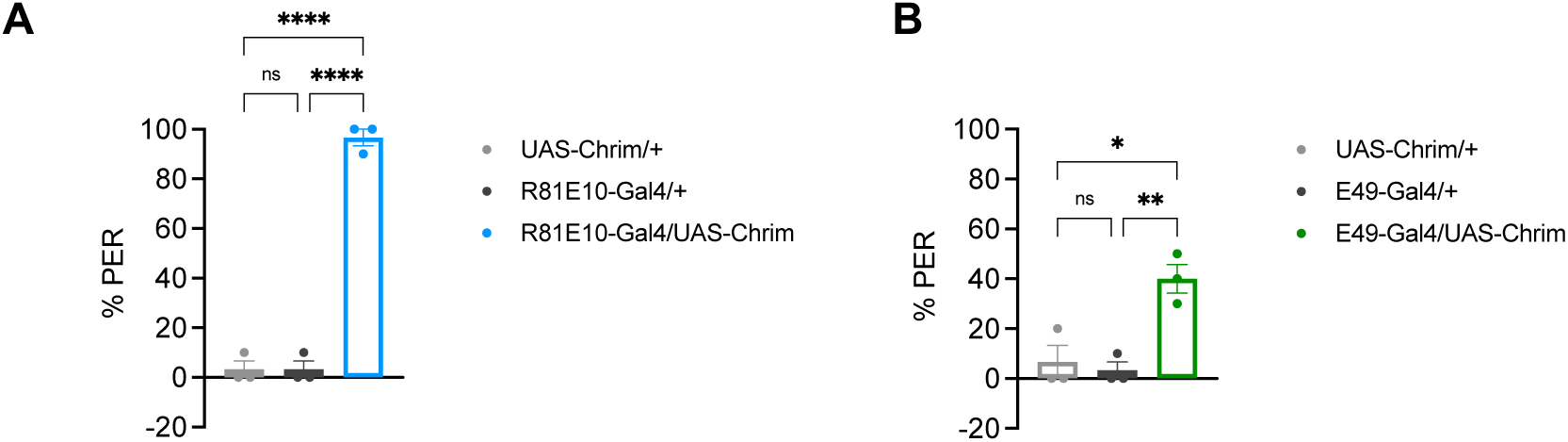
Activation of the Fdg neuron or MN9 motor neuron acutely elicits PER. (A-B) Percent of flies acutely showing PER during light activation of the Fdg neuron (A) or MN9 motor neuron (B) (n = 3 sets of flies). Genotypes were compared using one-way ANOVA followed by Tukey’s multiple comparisons test.

